# Rapidly Computing the Phylogenetic Transfer Index

**DOI:** 10.1101/743948

**Authors:** Jakub Truszkowski, Olivier Gascuel, Krister M. Swenson

## Abstract

Given trees *T* and *T*\* on the same taxon set, the *transfer index ϕ*(*b, T*\*) is the number of taxa that need to be ignored so that the bipartition induced by branch *b* in *T* is equal to some bipartition in *T*\*. Recently, Lemoine *et al*. [14] used the transfer index to design a novel bootstrap analysis technique that improves on Felsenstein’s bootstrap on large, noisy data sets. In this work, we propose an algorithm that computes the transfer index for all branches *b ∈ T* in *O*(*n* log^3^ *n*) time, which improves upon the current *O*(*n*^2^)-time algorithm by Lin, Rajan and Moret [15]. Our implementation is able to process pairs of trees with hundreds of thousands of taxa in minutes and considerably speeds up the method of Lemoine *et al*. on large data sets. We believe our algorithm can be useful for comparing large phylogenies, especially when some taxa are misplaced (e.g. due to horizontal gene transfer, recombination, or reconstruction errors).

## 1 Introduction

The need to compare phylogenetic trees arises in many contexts in computational biology. In bootstrap analysis [12], trees inferred from bootstrapped data sets are compared to the tree inferred from the original data set (known as the *reference tree*) in order to evaluate the statistical support for every branch in the inferred topology. Distance measures between phylogenies, such as the Robinson-Foulds distance [10] or quartet distance [7], are also frequently used to evaluate the accuracy of inference methods by comparing the trees they produce with the “ground truth” trees that were used to simulate the data set. Finally, distance measures between phylogenetic trees allow researchers to cluster trees inferred from different genes to reveal groups of genes with similar patterns of evolution due to horizontal gene transfer, duplication, loss, or recombination [13].

The most popular approach to comparing phylogenies compares the sets of splits (*bipartitions*) induced by the two trees. In the bootstrap setting, each branch in the reference tree is assigned a *bootstrap score* which is the fraction of bootstrap trees that contain a branch inducing the same split as the branch in the reference tree. Higher scores imply more confidence in the inferred branches. This is known as Felsenstein’s bootstrap (FBP). When the goal is to compare the global similarity of a pair of trees, a related approach is to compute the number of splits that are found in one of the trees but not both. This is known as the Robinson-Foulds(RF) distance and is widely used for comparing phylogenies. The RF distance can be computed in *O*(*n*) time thanks to an algorithm by Day [10].

Both FBP and the RF distance are widely used due to their low computational cost and ease of interpretation. However, these approaches become inadequate when dealing with large, noisy data sets where large splits are unlikely to be recovered exactly. For example, if the data set contains a small number of recombinant sequences, the location of those sequences may vary between the bootstrap trees and the reference tree. Even a small number of such sequences can impact many splits in the tree, leading to low bootstrap values across the tree, despite the fact that the location of most sequences is strongly supported by the data. This problem is exacerbated in large trees, where the risk of including a rogue taxon is elevated. To illustrate this problem, Lemoine *et al*. [14] showed that trees built on a random sampling of HIV strains have generally higher bootstrap values than a tree built on the full set of strains.

Recently, Lemoine *et al*. [14] addressed this shortcoming of FBP by introducing a new bootstrap procedure based on finding similar, rather than identical splits in the bootstrap trees. For each split *b* in the reference tree and each bootstrap tree *T*\*, their method computes the *transfer index ϕ*(*b, T*\*) between the split and the bootstrap tree, which is the minimum number of taxa that need to be removed from the taxon set for *b* to equal some split *b*\* in *T*\*. The bootstrap support value assigned to *b* is then 1.0 minus its transfer index normalized by the size (minus 1.0) of the smaller of the bipartition sets induced by *b*, averaged over all bootstrap trees (see Davila Felipe *et al*. [9] for the statistical justification of this formula). Lemoine *et al*. show that this approach is considerably more robust to the presence of unstable taxa. The transfer index is computed using a quadratic-time algorithm due to Lin, Rajan and Moret [15] (see also [3]), which is efficient for moderate-sized data sets, but becomes less practical for trees containing thousands or tens of thousands of taxa. In contrast, modern tree-building methods can reconstruct trees with hundreds of thousands of taxa [18, 5], which creates a need for scalable post-processing and analysis tools.

In this work, we present a fast exact algorithm for computing the transfer index for all edges in the reference tree with respect to a bootstrap tree. Our algorithm runs in *O*(*n* log^3^ *n*) time where *n* is the number of taxa in the data set, which is considerably faster than the *O*(*n*^2^) required by Lin, Rajan and Moret [15]. Our prototype Python implementation [1] is able to process pairs of trees with tens of thousands of taxa in a matter of seconds on a standard laptop computer. Our C implementation for balanced trees [2] enables us to carry out the analyses of Lemoine *et al*. at least an order of magnitude faster, and makes it possible to compute the transfer index on data sets with tens or hundreds of thousands of sequences. Computations that took nearly nine hours now take about ten minutes. Pairwise comparisons of trees with hundreds of thousands of taxa can be completed within minutes.

Simultaneously and independently of our work, Lutteropp, Kozlov and Sta matakis [16] developed an efficient implementation of the original algorithm that is up to 480 times faster than the original Booster implementation and requires 10x - 40x less memory in their experiments.

## 2 Preliminaries

We consider binary trees with *n* leaves uniquely labeled from a set *L* of size *n*. Take a tree *T*. A branch *b ∈ E*(*T*) can be described by the bipartition *{A, B}* that it induces on *L* when removed from the tree (*A ∪ B* = *L*). For *L*′ *⊆ L*, define *b | L*′ = *{A ∩ L*′, *B ∩L*′ *}*.

Consider two leaf-labeled binary trees *T* and *T*\* with branches *b ∈ E*(*T*) and *b*\**∈ E*(*T*\*). The *transfer distance δ*(*b, b*\*) is the number of leaves that must be ignored so that *b* and *b*\* are equal

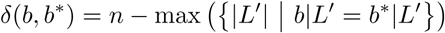

The *transfer index* for *b* with respect to *T*\* is the minimum transfer distance over all possible *b*\* *∈ E*(*T*\*):

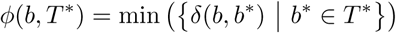

This work is about computing *ϕ*(*b, T*\*) for all *b ∈ E*(*T*). We first describe an algorithm for a special case of the problem, corresponding to situations where *T*\* is a balanced tree. We then present a modification of the algorithm that enables us to process general trees efficiently. Finally, we evaluate the performance of our algorithm on simulated and empirical data sets and discuss directions for future research.

## 3 An algorithm for balanced trees

In this section, we describe an efficient algorithm to compute *ϕ*(*b, T*\*) for all *b ∈ E*(*T*) when *T*\* is a balanced tree. For the purposes of this manuscript, we say that a tree *T*\* is *balanced* if there exists a node *r ∈ V* (*T*\*) (which we call the *root*) such that the path from *r* to any leaf of *T*\* contains at most *C* log *n* edges for some constant *C*. Under this assumption, our algorithm can compute *ϕ*(*b, T*\*) for all branches *b ∈ E*(*T*) in *O*(*n* log^3^ *n*) time. Balanced trees are common in phylogenetics, as both the Kingman coalescent (standard population genetics model) and Yule (standard speciation model) trees are expected to be balanced [20].

For convenience, we will work with transfer distances between rooted sub-trees. For a node *u ∈ V* (*T*), let *L*(*u*) be the set of leaves descendant from *u*. The *rooted transfer distance* is

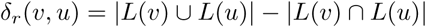

for *v ∈V* (*T*\*) and *u ∈ V* (*T*). The (unrooted) transfer distance between the corresponding splits *b*_*v*_ and *b*_*u*_ is then

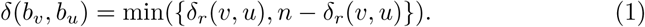

Let *T* be a rooted binary tree. For any node *u* in *V* (*T*), let *c*_*h*_(*u*) and *c*_*l*_(*u*) be the children of *u* such that *|L*(*c*_*h*_(*u*)) *| ≥| L*(*c*_*l*_(*u*)) *|*. We call *c*_*h*_(*u*) and *c*_*l*_(*u*) the *heavy* and *light* child of *u*, respectively. We refer to the edge (*u, c*_*h*_(*u*)) as the *heavy edge*. If both children of *u* have an equal number of leaf descendants, we pick the heavy child arbitrarily.

The main idea of the algorithm is to exploit the structure of both *T* and *T*\* when computing the transfer index for subsequent splits. Suppose that *T* contains two vertices *u, u* The main idea of the algorithm is to such that *L*(*u*^*’*^) = *L*(*u*) *∪{x}* for some leaf *x*. Unsurprisingly, we can show that the transfer distance values *δ*_*r*_(*v, u*) and *δ*_*r*_(*v, u*^*’*^) differ by at most 1.

### Lemma 3.1.

*Let u and u*^*’*^ *be nodes in V* (*T*) *such that L*(*u*^*’*^) = *L*(*u*) *∪{x} for some leaf x. Let P* = *v*_1_, *…, v*_*k*_ *be the path from x to r in T*\*, *where v*_1_ = *x and v*_*k*_ = *r. Then, for any* 1 *≤ i ≤ k*

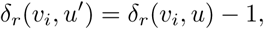

*and for any v ∉ P*

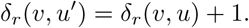

*Proof*. Any node *v* on the path between *r* and *x* in *T*\* has *x* as its descendant, so *δ*_*r*_(*v, u*′) = *δ*_*r*_(*v, u*) *−* 1 as *x* does not have to be removed from *L*(*v*) to obtain *L*(*u*^*’*^), but it has to be removed to obtain *L*(*u*). The opposite holds for nodes outside the path from *r* to *x*, so the distance increases by one – see Figure 1. □

**Figure 1:**
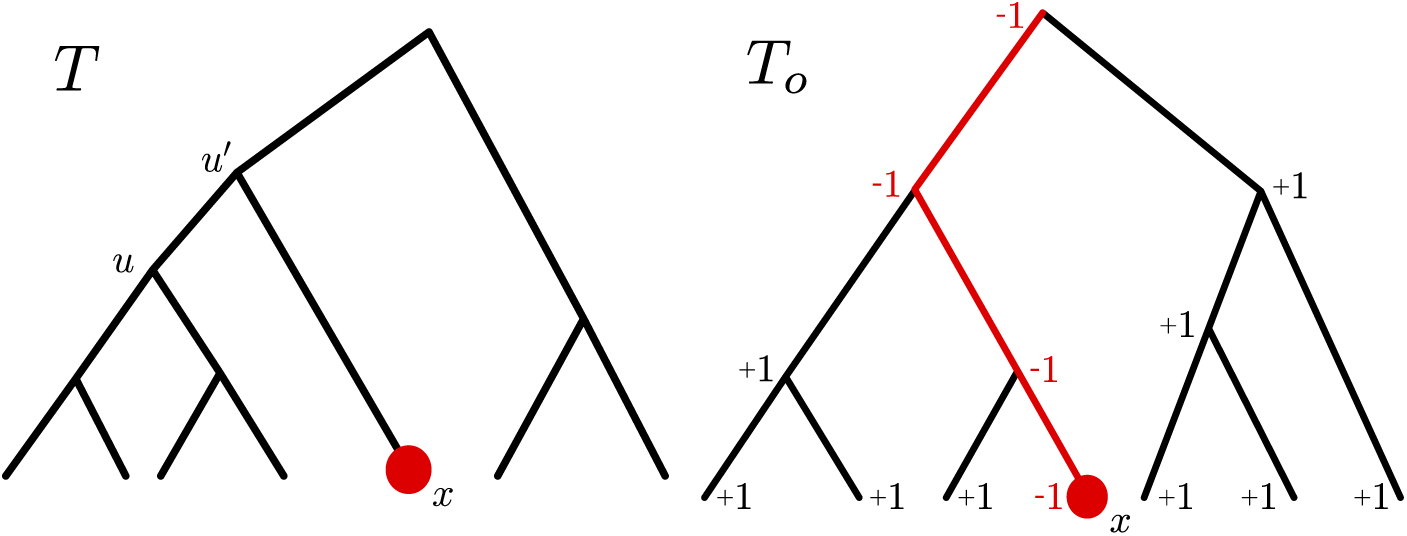
The difference between the rooted transfer distance to *u* and *u*′, for all nodes in *T*\*. The rooted transfer distance decreases for all nodes on the path from *x* to the root and increases for all other nodes.

The above lemma suggests a strategy for efficiently computing *δ*_*r*_(*v, u*′) from *δ*_*r*_(*v, u*). For each node *v ∈ V* (*T*\*), we will maintain a variable *D*[*v*] which, when combined with a global counter, maintains the invariant *δ*_*r*_(*v, u*) = *D*[*v*] + *counter*. Algorithm *AddLeaf* updates *D* when moving from *u* to *u*′. The global counter is incremented, effectively increasing the transfer distance for all nodes. To compensate for this, each *D* along the path from *x* to the root is decreased by 2, thus maintaining the invariant.

### Algorithm 1 AddLeaf(*x*)

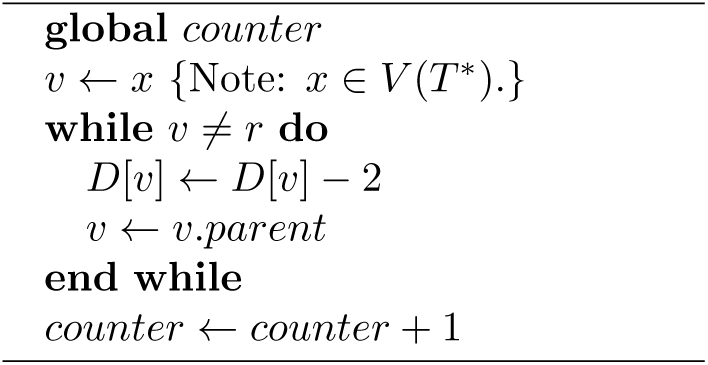

### Algorithm 2 RemoveLeaf(*x*)

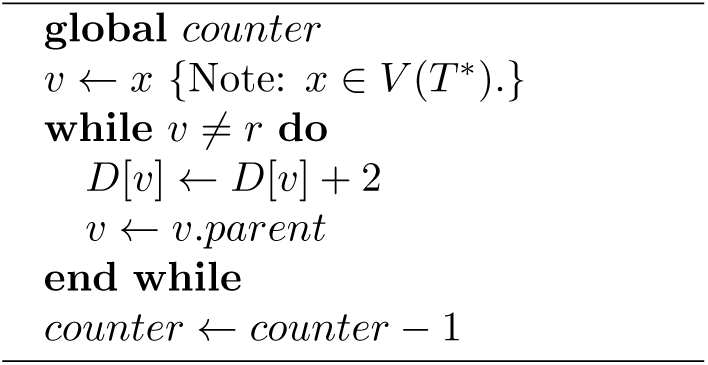

Finding the node *v* that minimizes *δ*_*r*_(*v, u*) can be achieved by using a dynamic data structure similar to a heap [8]. The total running time of *AddLeaf* is *O*(log^2^ *n*) for balanced trees as the main loop is executed *O*(log *n*) times and each update of the structure takes *O*(log *n*) time.

When *L*(*u*′) differs from *L*(*u*) by more than one leaf, we update *D* by calling *AddLeaf* for all leaves in *L*(*u*′) *− L*(*u*).

We now present the complete algorithm as Algorithm 3. First, we initialize the *D* values by computing the rooted transfer distance from the empty subtree to every subtree in *T*\*. The rooted distance from the empty subtree equals the number of leaves in *L*(*v*). Then, we choose a leaf *x* in *T* and traverse its ancestors, calling *AddLeaf* on leaves descended from nodes off the path from *x* to *r*. To ensure that *AddLeaf* is called a limited number of times on a leaf, we terminate the traversal when the current node is the light child of its parent. In that case, we undo the leaf additions using the routine *RemoveLeaf* and restart the process from a new leaf.

### Algorithm 3 ComputeTransferIndices(*T*\*, *T*)

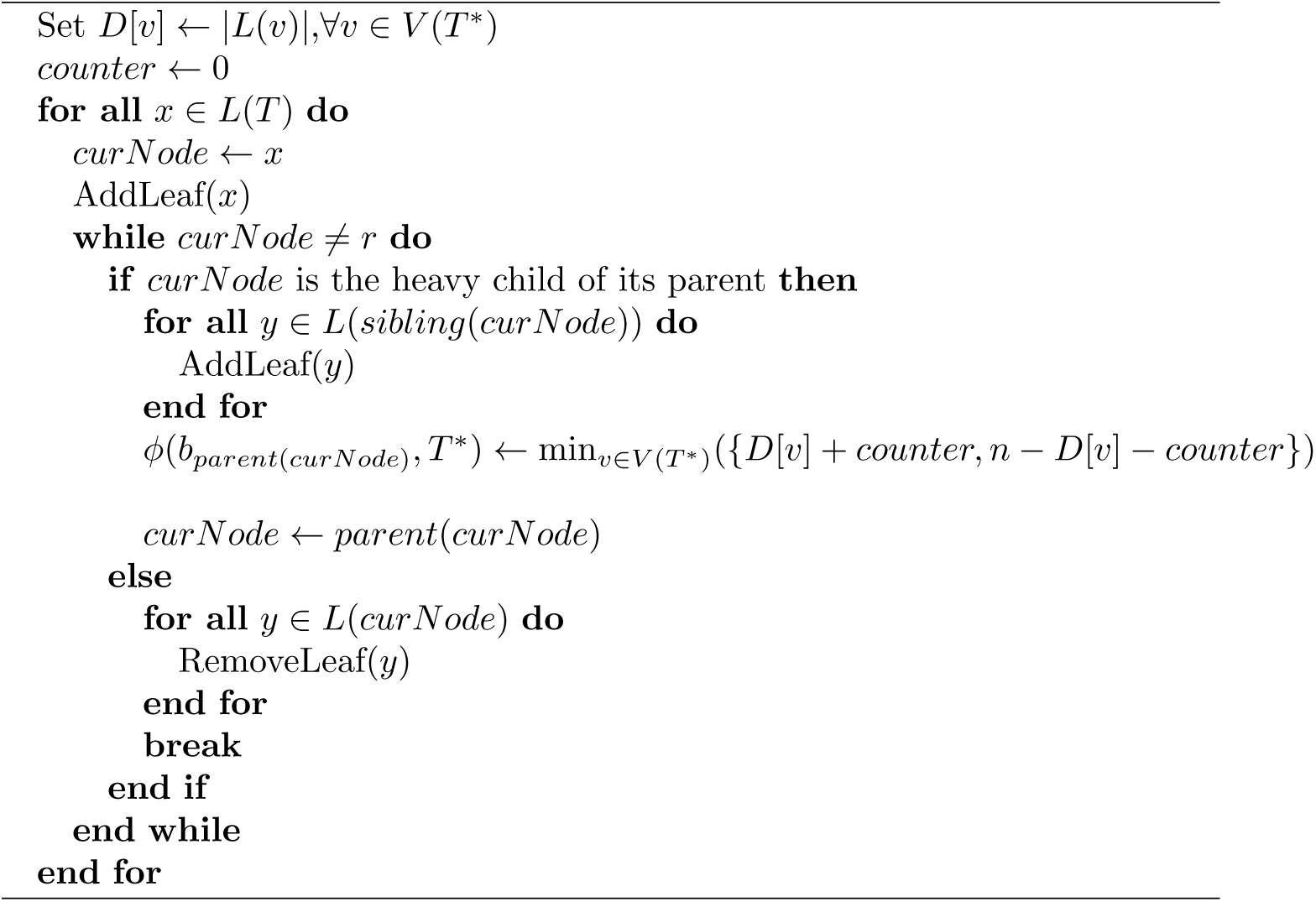

We can prove the following property.

### Lemma 3.2.

*The number of calls made to AddLeaf during the execution of ComputeTransferIndices is at most n* log2 *n*.

*Proof*. Let *x* be a leaf in *T*. The number of times *AddLeaf* is called with *x* given as the argument is at most the number of nodes *v* on the path from *x* to the root such that 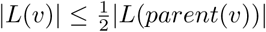. There can be at most log2 *n* such nodes since *|L*(*r*) *|*= *n* and *|L*(*x*) *|* = 1. We obtain the result by summing over all leaves in *T*. □

Since the running time of *AddLeaf* is *O*(log^2^ *n*), it follows that the total running time of the algorithm is *O*(*n* log^3^ *n*).

## 4 General Trees

### 4.1 The data structure

The performance of Algorithm 3 crucially depends on the height of the phylogeny. When *T*\* is a caterpillar tree and *T* is balanced, the running time of the algorithm can be as bad as *O*(*n*^2^ log^2^ *n*). In order to derive a fast algorithm for general trees, we need a data structure that would allow us to efficiently navigate splits in the tree, regardless of the topology. Our approach is based on *heavy path decompositions*, originally introduced by Sleator and Tarjan [19].

Recall that a heavy child is a node that has at least as many descendant leaves as its sibling, and that a heavy edge is an edge that connects a heavy child to its parent. A *heavy path* is a sequence of nodes *v*_1_, *…, v*_*k*_ such that (*v*_*i*_, *v*_*i*+1_) is a heavy edge for all 1 *≤ i ≤ k −* 1 (see Figure 2). A heavy path is called *maximal* if *v*_*k*_ is a leaf and *v*_1_ is either the root or the light child of its parent. We let 𝒫(*T*) denote the set of maximal heavy paths in *T*.

**Figure 2:**
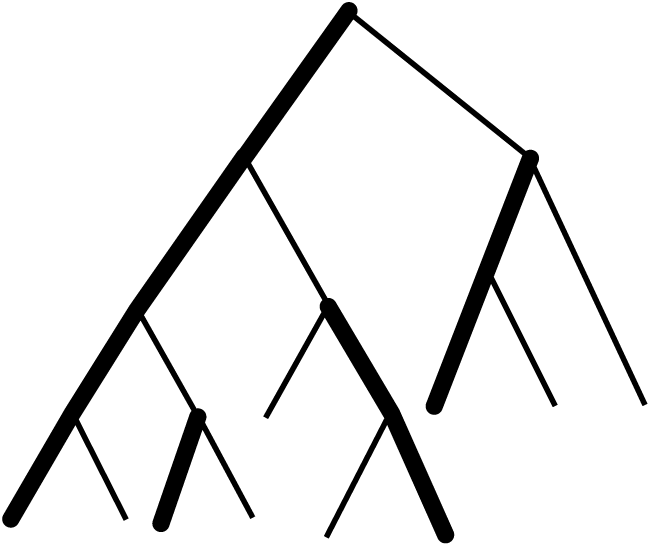
A phylogeny with heavy paths marked in thick lines.

#### Lemma 4.1.

*Let x be any leaf in T*\*. *The path from x to the root of T*\* *intersects at most j*log *nl* + 1 *distinct maximal heavy paths*.

*Proof*. Let (*u, v*) be an edge on the path from *x* to the root such that *u* and *v* belong to different maximal heavy paths. Without loss of generality, we can assume that *u* is the parent of *v*. Then *|L*(*u*) *| ≥*2 *|L*(*v*) *|* since otherwise (*u, v*) would be a heavy edge. Since *L*(*x*) = 1 and *L*(*root*(*T*\*)) = *n*, it follows that there are at most *j*log *nl* such edges, which gives the result. □

We will think of *T*\* as a collection of maximal heavy paths. For each maximal heavy path *p* = *v*_1_, *…, v*_*k*_, we recursively define a *path search tree* 𝒮(*p*) as follows. The root of 𝒮(*p*) is associated with *p*. The children of a node associated with a path *p*′ = *v*_*p*′_1, *…, v*_*p*′_*k′* are associated with subpaths *v*_*p*′_1, *…, v*_*p*′_⌊*k′/*2⌋ and *v*_*p*′_⌊*k’/*2+1⌋, …, *v*_*p*′_*k*′; we refer to them as *intervals* and write [*a, b*] to denote both the node associated with the path *a, …, b* and the path itself. If *p*′ has only one element, it is a leaf in 𝒮 (*p*). We also write *first*(*p*) := *v*_1_, *last*(*p*) := *v*_*k*_ and *path*(*x*) for the unique maximal heavy path that contains *x*.

For each interval [*a, b*] we maintain variable *D*[*a, b*] and maintain the invariant

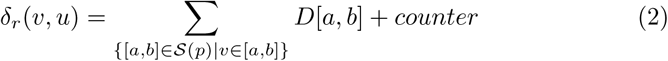

for all *v ∈ V* (*T*\*), where *u* is the node in *T* under consideration. Moreover, we maintain variables *minval*[*a, b*] and *maxval*[*a, b*] with the invariant

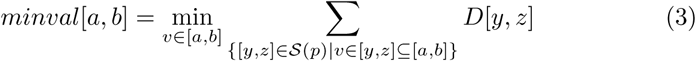

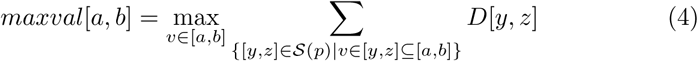

Informally, *minval*[*a, b*] is the minimum value of the rooted transfer distance between any node in [*a, b*] and the current split in *T*, up to a constant. That is, the node *v* that minimizes ∑_{[*y,z*]∈𝒮(*p*)*|v∈*[*y,z*]*⊆*[*a,b*]}_ *D*[*y, z*] also minimizes *δ*_*r*_(*v, u*). In particular, if [*a, b*] is the root node of its path search tree, we have

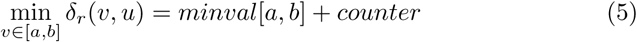

Analogously, for *maxval*[*a, b*] we have

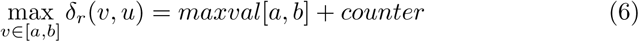

While we are not aware of this particular data structure having been defined previously, we note that somewhat similar search tree structures on phylogenies have been developed before for large-scale phylogenetic tree reconstruction [6] and computing the quartet distance between trees [4]. Heavy path decompositions have also been introduced in database literature for computing edit distances on trees [17].

### 4.2 The algorithm

Algorithm 7 follows the same strategy as the algorithm for balanced trees, with a few modifications designed to accommodate potentially long paths between the root and leaves in *T*\*. First, we initialize the path search trees by setting *D*[*x, x*] = *|L*(*x*) *|* for every node *x ∈ V* (*T*\*) and *D*[*a, b*] = 0 for every interval with *a* ≠ *b*. We also set *minval*[*a, b*] = *|L*(*b*) *|*, *maxval*[*a, b*] = *| L*(*a*) *|*, and *counter* = 0. It can be easily verified that the invariants given by Equations 2, 3, and 4 are satisfied.

The algorithm then repeatedly calls functions *AddLeaf General* and *RemoveLeaf General* to move between splits of *T*, updating the transfer distances to splits in *T*\*. We use path search trees together with the function *UpdatePath* to efficiently update the minimum transfer distance values within each heavy path, making use of Equations 5 and 6. Given a node *x*, the algorithm traverses the intervals in the path search tree of the heavy path containing *x* while updating *D, minval*, and *maxval* values to reflect the fact that the distance has changed by the same amount for *x* and every node above it (see Figure 3).

**Figure 3:**
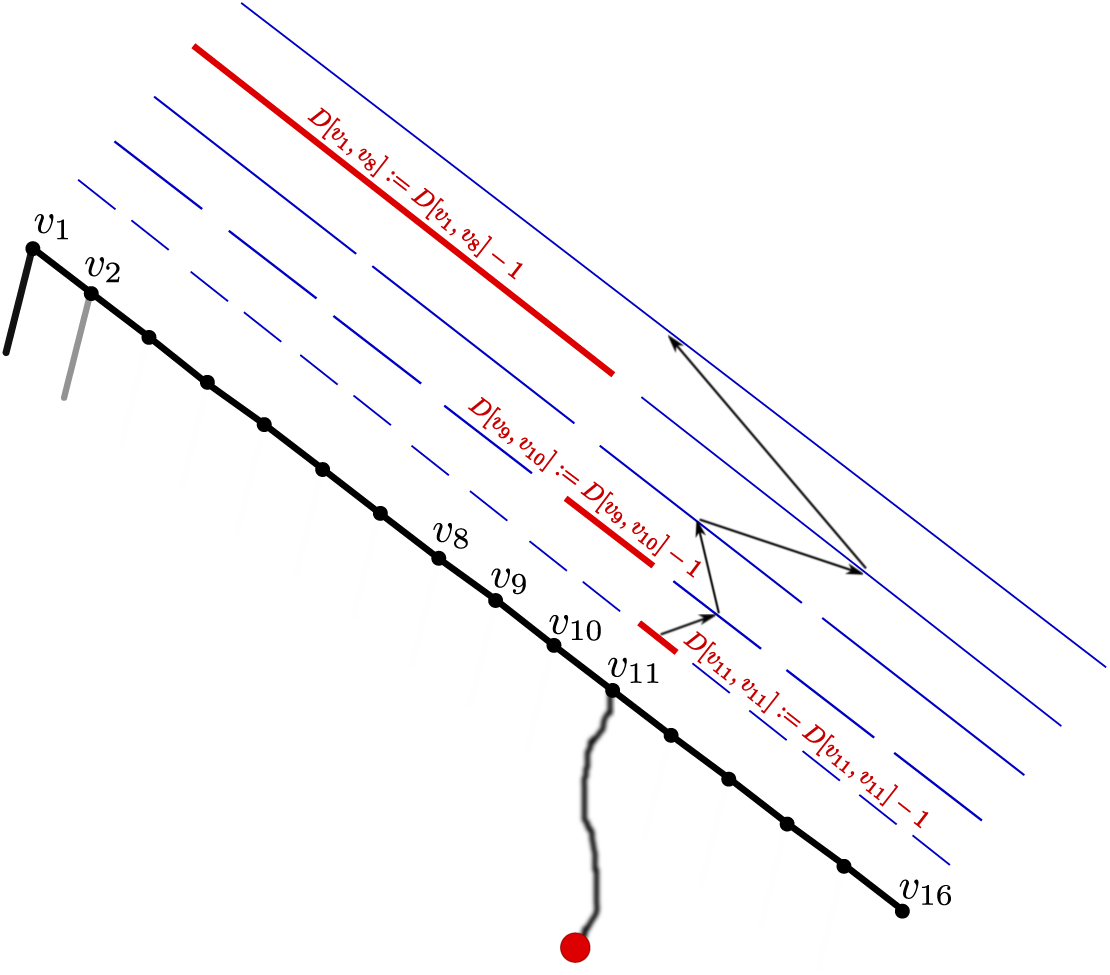
Updating the *D* values associated with the heavy path *v*_1_, *…, v*_16_ during the call to *UpdatePath*(*v*_11_, *−* 2). Intervals whose *D* values have been updated are coloured in red.

#### Algorithm 4 UpdatePath(*x,d*)

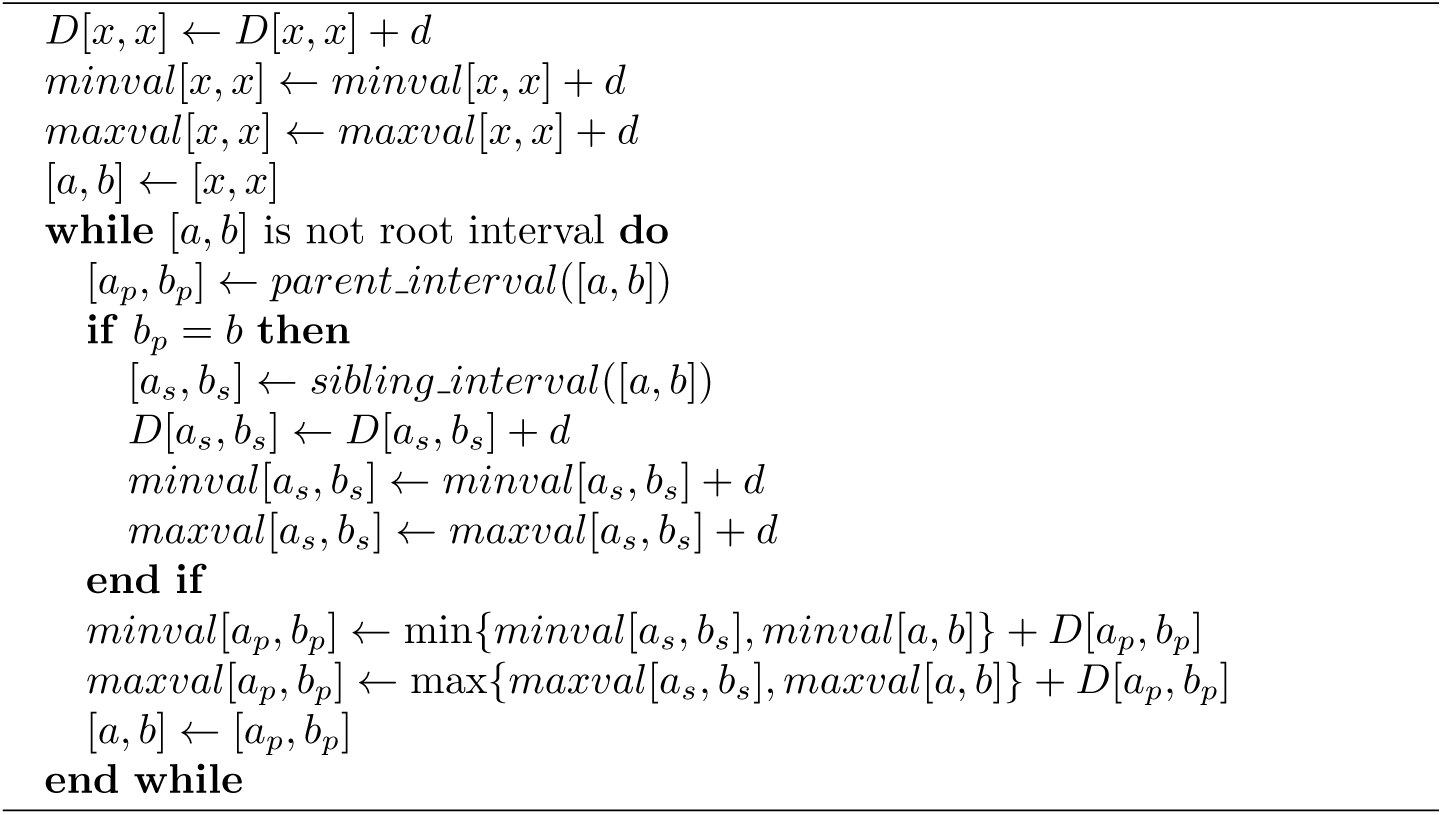

#### Lemma 4.2.

*Each call to UpdatePath changes the value of ∑* _{[*a,b*]*∈𝒮*(*p*)*|v∈*[*a,b*]*}*_ *D*[*a, b*] *by d for all vertices v ancestral to x (including x) in the path p. Moreover, each call preserves invariants 3 and 4*.

*Proof*. Let *y* be a node in *p* that is above *x*. Let [*a*_*m*_, *b*_*m*_] be the minimal interval in 𝒮(*p*) that contains both *x* and *y* and let [*a*_*l*_, *b*_*l*_] and [*a*_*r*_, *b*_*r*_] be the left and right child of [*a*_*m*_, *b*_*m*_] in 𝒮 (*p*), respectively. By the choice of [*a*_*m*_, *b*_*m*_], *x* and *y* cannot both belong to the same child of [*a*_*m*_, *b*_*m*_]. Since *y* is above *x* in *T*\*, it follows that *y ∈* [*a*_*l*_, *b*_*l*_] and *x ∈* [*a*_*r*_, *b*_*r*_]. We have *b*_*m*_ = *b*_*r*_ by the definition of the path search tree, so *D*[*a*_*l*_, *b*_*l*_] will be increased by *d* at the iteration of the *while* loop when [*a, b*] = [*a*_*r*_, *b*_*r*_] and *minval*[*a*_*l*_, *b*_*l*_] and *maxval*[*a*_*l*_, *b*_*l*_] will be increased by the same amount, preserving invariants 3, 4. On the other hand, it can be easily verified that all the other variables *D*[*a, b*] such that *y ∈* [*a, b*] will remain unchanged, since [*a*_*l*_, *b*_*l*_] is the only interval containing *y* that is the left child of its parent and has a right child that contains *x*. □

#### Algorithm 5 AddLeafGeneral(*x*)

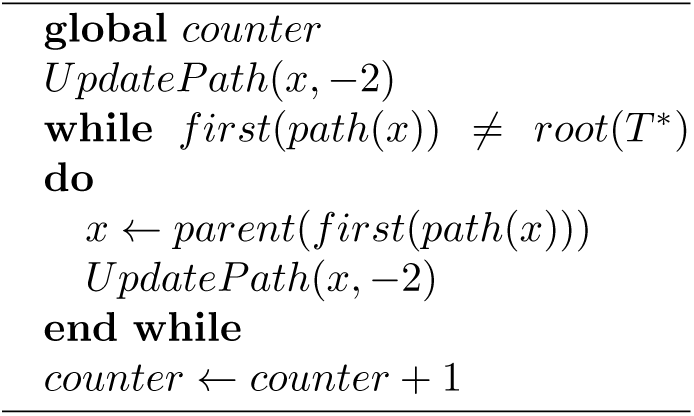

#### Algorithm 6 RemoveLeafGeneral(*x*)

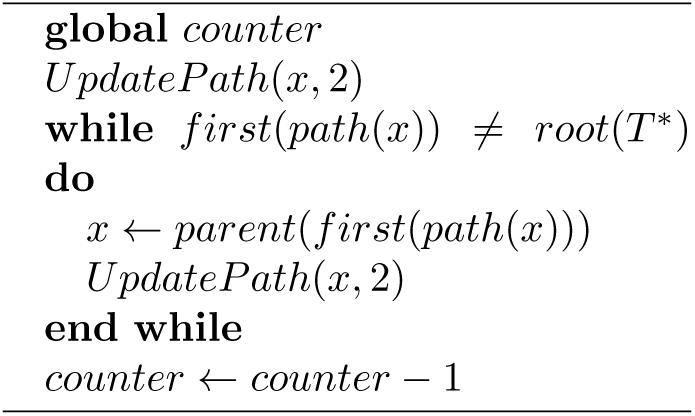

#### Lemma 4.3.

*After each call to AddLeaf General or RemoveLeaf General, Equation 2 is satisfied*.

*Proof*. Equation 2 holds for the initial values of *D*[*a, b*] and *counter*. The calls to *UpdatePath* from within *AddLeaf General* decrease the sum ∑ _*{* [*a,b*] *∈*𝒮 (*p*) *| v ∈* [*a,b*]}_ *D*[*a, b*] by exactly 2 for each node *v* on the path from *x* to the root of *T*\*. The variable *counter* increases by 1, which means that the sum

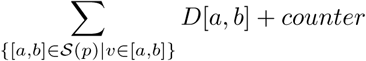

is decreased by 1 for each node on the path from *x* to the root and increased by 1 for all other nodes. This, together with Lemma 3.1 gives the result for *AddLeaf General*. The case of *RemoveLeaf General* is proved by analogous reasoning. □

#### Theorem 4.1.

*Algorithm 7 computes the transfer index for all splits in T*.

*Proof*. Lemma 4.3 and Equation 5 ensure that at each iteration of the *while* loop, the value *minval*[*first*(*p*), *last*(*p*)] + *counter* is equal to the minimum value of the rooted transfer distance among the nodes in *p*. Using Equation 1 and taking the minimum over all the paths yields the result.□

#### Algorithm 7 ComputeTransferIndicesGeneral(*T*\*, *T*)

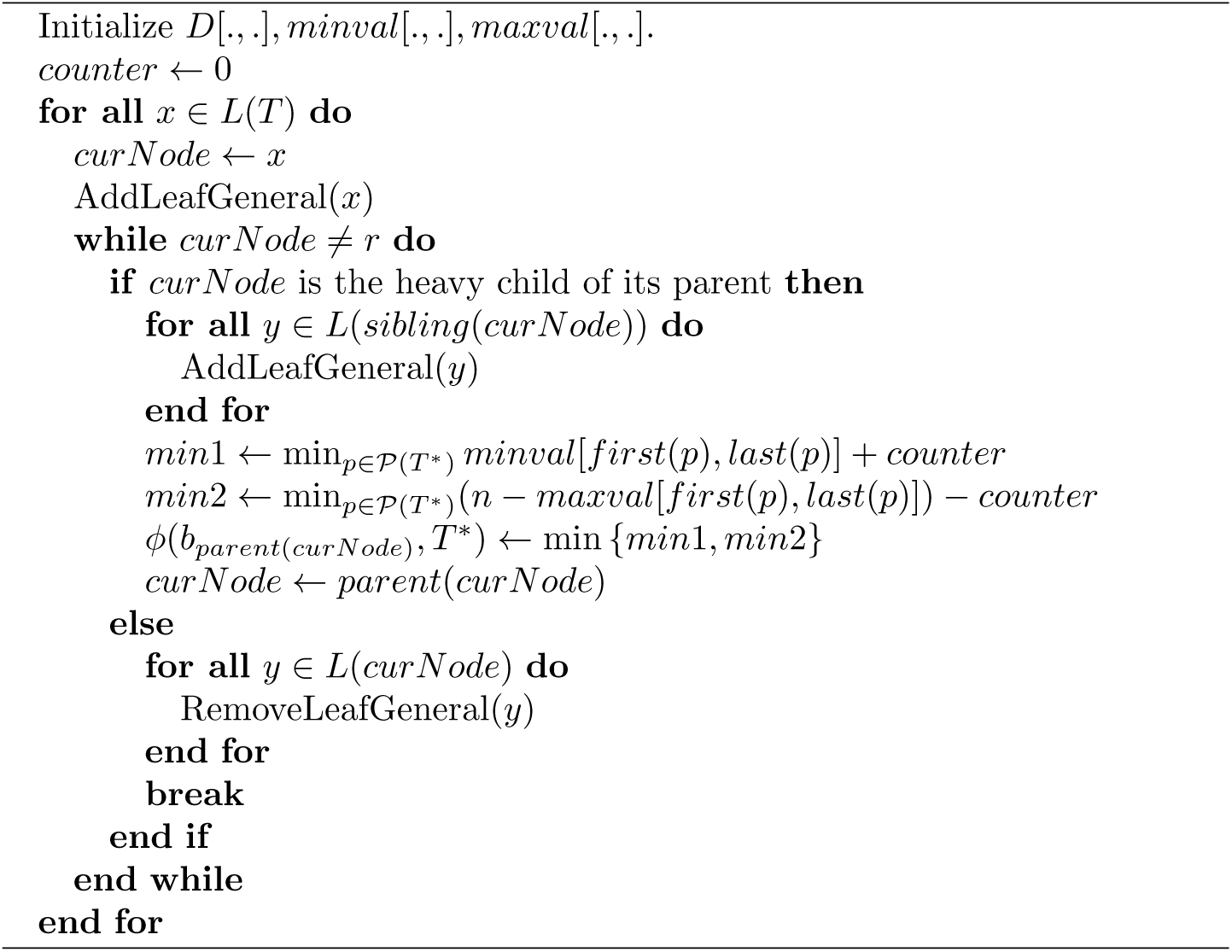

### 4.3 Running time analysis

The running time of Algorithm 7 is *O*(*n* log^3^ *n*). Each call to *UpdatePath* takes at most *O*(log *n*) time. During the execution of *AddLeaf General* and *RemoveLeaf General, UpdatePath* is called at most *O*(log *n*) times by Lemma 4.1, which gives a bound of *O*(log^2^ *n*) for each call to any of these functions. Finally, by Lemma 3.2, *AddLeaf General* is executed at most *O*(*n* log *n*) times, which means that the overall running time is *O*(*n* log^3^ *n*). Finding *ϕ*(*b*_*curNode*_, *T*\*) can be solved efficiently using a heap at a cost of *O*(log *n*) for each call to *UpdatePath*, which increases the total running time only by a constant factor.

## 5 Implementations

We have two implementations currently available. A proof-of-concept implementation of Algorithm 7 was written in Python, making use of the Dendropy package [21]. This implementation is available at [1]. A definitive C implementation is a work in progress; Algorithm 3 has been integrated into the current Booster code base [2].

To obtain speedups for trees with a high degree of similarity, our Python implementation first traverses both trees and identifies, in linear time, maximal identical subtrees that occur in both trees. These trees are then replaced with single nodes whose weight equals the number of leaves in the collapsed subtree. After this pre-processing step, every *AddLeaf* and *RemoveLeaf* operation changes all the relevant counters by the weight of the node, rather than by 1. Every edge within the collapsed subtrees has transfer index equal to 0. This procedure can result in substantial speedups for highly similar trees (see next section).

## 6 Experiments

We evaluated the performance of our algorithm on several simulated and empirical data sets. In all simulation experiments, we simulated pairs of trees by sampling the first tree topology uniformly at random and applying random edit operations to obtain the second tree. Each edit operation chose two random nodes such that neither node is ancestral to the other (with respect to some arbitrary rooting of the tree) and swapped the subtrees rooted at the chosen nodes. By increasing the number of applications of the edit operation, we decreased the similarity of the trees.

In the first experiment, we generated pairs of trees with sizes varying between 100 and 100000 taxa. Each pair of trees was generated using the number of edit operations equal to 0.2 times the number of taxa. Experiments were performed on a Linux laptop with 8 GB of RAM and an Intel i5 2.3 GHz processor using one thread for each run. The results are given in Figure 4(left). The running time of our Python program is close to linear in the number of taxa; even for 100000 taxa, it takes less than 100 seconds to compute the transfer indices. Our C implementation of Algorithm 3 is faster for smaller trees with up to 20000 taxa, but becomes considerably slower for very large trees. This is not surprising as trees sampled from the uniform distribution are likely to be unbalanced, as the mean diameter of a tree is 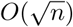 [11]. Booster is the slowest of the three algorithms and runs out of memory for trees with more than 20000 taxa.

**Figure 4:**
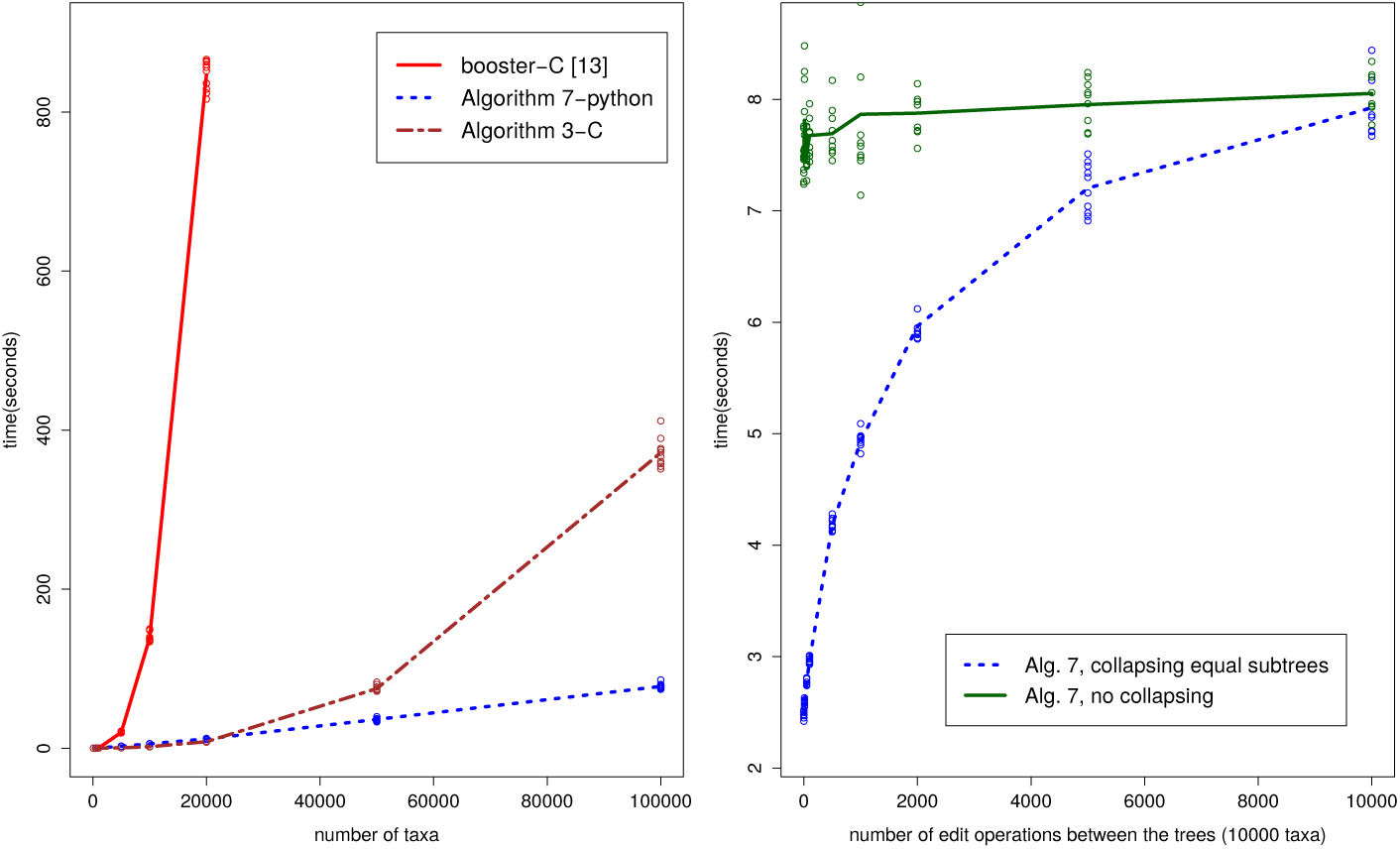
Left: The running times for our two implementations and the code by Lemoine *et al*. as a function of the number of taxa. Right: Running times for a pair of 10000-taxon trees as a function of the similarity of the trees.

In the second experiment, we investigated the impact of tree dissimilarity on the running time. Experiments were performed on an Intel Xeon 2.1 GHz processor with 32 GB of RAM using a single thread. We generated pairs of trees with 10000 taxa each while varying the number of edit operations from 0 to 10000. As we can see in Figure 4(right), tree dissimilarity has little, if any, effect on the running time of the algorithm when the preprocessing procedure is not applied. In contrast, collapsing identical subtrees can lead to 2- to 3-fold speedups for highly similar tree pairs.

In the final experiment, we repeated the analysis from Lemoine *et al*. on several HIV and mammalian data sets. Each data set contained 1000 bootstrap replicates and the number of taxa varied between 571 and 9147. The results are given in Table 1. On the largest data set, our Python program completed the analysis in less than 2 hours and our C program completed the analysis in less than 12 minutes, compared to almost 9 hours for Booster, which is written in C. Smaller data sets were processed in a matter of minutes by Booster and our Python implementation, while our C implementation was at least an order of magnitude faster than Booster in all cases.

**Table 1:** Running times for four data sets from Lemoine *et al*.. Our C implementation presents substantial advantages.

| Data set | taxa | #bootstraps | booster-C [14] | Alg. 7-Python | Alg. 3-C |
| --- | --- | --- | --- | --- | --- |
| HIV/Full | 9147 | 1000 | 8h51m | 1h47m | **12m** |
| HIV/Medium_Sample1 | 571 | 1000 | 1m | 3m23s | **6s** |
| HIV/Medium_Sample2 | 571 | 1000 | 1m | 3m32s | **6s** |
| Mammals/raxml | 1449 | 1000 | 10m32s | 10m36s | **30s** |

## 7 Conclusion

We have presented an algorithm for rapidly computing, for every split in tree *T*, the minimum transfer distance to the closest split in another tree *T*\*. The running time of the algorithm is *O*(*n* log^3^ *n*), which is close to linear in the size of input trees. We expect to be able to remove a logarithmic factor with improved bookkeeping to maintain the minimum value of the transfer distance. Without this improvement, our prototype implementation scales almost linearly in practice, enabling us to compare pairs of trees with hundreds of thousands of taxa in a matter of minutes. This is comparable to several widely-used implementations of the RF distance, which makes our tool an interesting alternative to the RF distance for comparing large, noisy trees.

The most direct practical implication of this work is scaling up the TBE bootstrap analysis by Lemoine *et al*. Our method gives at least an order of magnitude speedup on all of the data sets they analyzed, and will enable the analysis of bootstrap data sets for trees of tens to hundreds of thousands of taxa within hours of CPU time.

On small and moderate-size data sets, our Python implementation is slower than the Booster software of Lemoine *et al*., which is likely due to their optimized implementation. Our C implementation for balanced trees always significantly out-performs Booster, however, and we are in the process of expanding it to the complete Algorithm 7.

There are several possible directions for future research. It would be interesting to see if the techniques developed here could be used to design novel methods for finding consensus trees. Another natural direction is to investigate whether global distance measures between trees, rather than splits, could be designed based on the transfer index.

## 8 Acknowledgements

We would like to thank Jeet Sukuraman for his help with the Dendropy package.

